# Nitrogen addition pulse has minimal effect in big sagebrush *(Artemisia tridentata)* communities on the Pinedale Anticline, Wyoming (USA)

**DOI:** 10.1101/446294

**Authors:** Christopher W. Beltz, Megan L. Mobley, Ingrid C. Burke

## Abstract

Nitrogen additions are known to elicit variable responses in semi-arid ecosystems, with responses increasing with precipitation. The response of semi-arid ecosystems to nitrogen are important to understand due to their large spatial extent worldwide and the global trend of increasingly available nitrogen. In this study, we evaluated the impact of a single nitrogen addition pulse on a semi-arid big sagebrush (*Artemisia tridentata*) ecosystem in western Wyoming. This is important given that sagebrush ecosystems are poorly understood, despite their prevalence in the western US. In addition, large-scale nitrogen additions have begun on sagebrush landscapes in Wyoming in order to mitigate population declines in mule deer (*Odocoileus hemionus*). The study objectives were (1) to evaluate the effectiveness of a nitrogen fertilization pulse in increasing sagebrush biomass and forage quality, and (2) to assess effects of nitrogen addition on soil biogeochemistry and vegetation community structure. We fertilized 15 plots across 5 locations in western Wyoming using a single pulse of urea (5.5g N m^−2^). In addition, we immobilized available nitrogen through surface hay treatments (250g hay/m^2^). Nitrogen additions failed to increase growth of sagebrush, alter nitrogen content of sagebrush leaders, or alter greenhouse gas efflux from soils. The plant community also remained unchanged; total cover, species richness, and community composition were all unaffected by our treatment application. Over the two years of this study, we did not find indications of nitrogen limitation of ecosystem processes, despite a wet growing season in 2014. Thus, we have found a general lack of response to nitrogen in sagebrush ecosystems and no treatment effect of a single pulse of N to sagebrush biomass or forage quality.

## INTRODUCTION

Nitrogen (N) additions are known to elicit varying responses in semi-arid ecosystems (1). Ecosystems respond to N in multiple ways, including increased net primary production and plant growth (2,3), plant community composition changes (4), and increased greenhouse gas (GHG) emissions from soils (5). Most semi-arid and arid systems (henceforth ‘drylands’) are thought to be primarily limited by water, though this limitation can be affected by both intra- and inter-annual variation in precipitation (6). If drylands were only restricted by water-limitation, N additions to the system should have no effect without a concurrent increase in precipitation. However, most dryland systems show a response to N additions alone, indicating at least a partial limitation by N (1,7).

In a meta-analysis of research studies on the effects of N fertilization studies in water-limited drylands worldwide, Yahdjian et al. (8) found that in almost all cases the addition of N increased net primary productivity (NPP), showing that nitrogen limitation – even if minor – is a global phenomenon. The ability to predict the effects of increased N on ecosystem process and structure is important because of the global trend of increasing N deposition (9). Given that drylands cover 40% of the Earth’s terrestrial surface (10,11) and 20% of the US (12), it is critical to investigate the effects of increasing N availability in dryland systems, due to their co-limitation by both water and N. In addition, dryland systems are home to 44% of all cultivated land (13,14), making understanding the effects of increasing N particularly important due to the high use of N-based fertilizers.

In the western US, big sagebrush (*Artemisia tridentata* spp.) ecosystems are the dominant dryland system (15,16), covering 48 million hectares (ha) and 13 states (17). Despite its current extent in the US, up to 60% of the historic, pre-settlement big sagebrush range has been partially or completely converted to exotic annual grasslands because of increased disturbance to these systems and conversion to grazing and agricultural lands (18). In addition, significant portions of the big sagebrush range continue to be affected by energy development, agriculture, and expansion of urban areas (19–22). These drivers of the loss of big sagebrush — unsustainable use of natural resources, population expansion, globalization, as well as climate change — are the same as those exacerbating desertification in drylands globally (13). Big sagebrush ecosystems are expected to continue to decline in coverage over the next decade with 2.3-5.5 million hectares projected to be impacted due to oil and gas development (23). While agricultural additions of N account for a majority of the increase in N availability globally, energy development also affects N deposition locally, through the production and release of NO_x_ compounds (9,24,25).

The effects of nitrogen additions on big sagebrush ecosystems are poorly understood due to both the small number of studies and the complexities associated with potential co-limitation of ecosystem processes by water and N in dryland systems (26). Although current knowledge gaps exist regarding N limitation in dryland ecosystems, increased N availability is widely known to have environmental consequences in other ecosystems (5). These include: (1) increasing the amount of carbon stored in terrestrial biomass in both soil and aboveground plant biomass (27,28), (2) increasing soil efflux of GHGs (29), and (3) changing the species composition of plant communities (30). Over the past decade, it has become increasingly important to understand the role of nitrogen in sagebrush ecosystems. Our research will help to close the gap in knowledge on the effects of N fertilization in sagebrush.

### Mule Deer: A Case Study in the Nitrogen Limitation of Drylands

The Pinedale Anticline of western Wyoming is one of the most disturbed regions of big sagebrush. Land managers are particularly concerned with large population declines in mule deer (*Odocoileus hemionus*) over the 15 years since the onset of natural gas development (31). The direct loss of 615 ha of big sagebrush habitat, and the associated winter forage, appear to be associated with the increased mortality in the Sublette Mule Deer Herd (32). By 2010, this population decline had been of sufficient magnitude that the Pinedale Anticline Project Office (PAPO) was required to initiate on-site mitigation.(26). Land managers hypothesized that fertilizing the area with N would increase habitat quality (33) through increases in big sagebrush biomass (i.e., forage production) and N content within the forage (i.e., forage quality).

The Pinedale Anticline is over 12,000 ha, with 320 ha in the pilot fertilization area (26,32). A single pulse of N fertilizer was applied by helicopter in 2010 and 2011. In 2010, two 100-acre plots were fertilized at 45 and 90 kg/ha. In 2011, a separate 400-acre plot was fertilized at the lower of the two rates (45 kg/ha). Each fertilization application was projected to cost US$135/ha – totaling greater than US$1.6m per year. Due to the high cost, potential environmental effects, and the expected dominance of water limitation, some ecologists argued that fertilization of big sagebrush was unlikely to be effective in offsetting forage loss for mule deer (26). The eventual monitoring report of the fertilization project reached a similar conclusion (34), however, as the study was conducted on a previously applied and un-replicated treatment, no generalizable conclusions could be reached.

N availability is likely to increase through the end of the century, largely due to the chronic addition of N to environments through N deposition (9). The chronic application (i.e., “press”) of N is likely to yield different effects than a single pulse, which we applied in this study to mimic the PAPO protocol (35,36). However, the development of a N fertilization project on the Pinedale Anticline provides a rare opportunity to ask basic research questions about N limitation in drylands, while simultaneously helping to inform decisions about a regional environmental issue.

### Project Goals & Hypotheses

Inspired by the actions on the Pinedale Anticline, the goals of this study were (1) to evaluate the efficacy of the addition of a N pulse for increasing big sagebrush biomass and forage quality, and (2) to assess effects of N addition on soil biogeochemistry, plant biomass, and plant community composition. We hypothesized that the N fertilization pulse would increase N availability in the soil, however it would not increase growth, biomass, or forage quality of big sagebrush due to primary limitation by water. In addition, we expected changes in plant community composition and a high potential for increased GHG emission.

## MATERIALS AND METHODS

### Study area and site selection

We conducted this study between 2012 and 2015 in semi-arid big sagebrush plant communities in western Wyoming, near the town of Pinedale (42°51′58″ N 109°51′53″ W). The area surrounding Pinedale is consistent with sagebrush steppe in other parts of Wyoming; plant communities are co-dominated by Wyoming big sagebrush (*Artemisia tridentata wyomingensis*) and mountain big sagebrush (*Artemisia tridentata vaseyana*). Other common plant species present include Sandberg bluegrass (*Poa secunda*), western wheatgrass (*Pascopyrum smithii*), chaffweed (*Antennaria microphylla*), squirreltail (*Elymus elymoides*), and longleaf phlox (*Phlox longifolia*).

The soils in this area are categorized as Typic Haplocryalfs, Typic Dystrocryepts, and Typic Haplocryolls with aridic moisture regimes (37). Soil textures at our sites were categorized as loam, sandy loam, and sandy clay loam (Table 1). Mean annual precipitation across the region averages 290 mm and mean annual temperature is 2.6 °C (Table 2) (38). The precipitation in this region falls predominantly in the spring and summer months.

**Table 1.** Soil texture classes and sand/silt/clay content for each exclosure. Ranches are the first letter in the ID (i.e., B, P, or S), site is the number in the second position, and the third letter corresponds to an individual plot (i.e., A, B, or C).

| Plot ID | Sand (%) | Clay (%) | Silt (%) | Soil Class |
| --- | --- | --- | --- | --- |
| 1-1A | 26 | 24 | 49 | loam |
| 1-1B | 43 | 24 | 32 | loam |
| 1-1C | 43 | 16 | 40 | loam |
| 1-2A | 41 | 16 | 42 | loam |
| 1-2B | 47 | 14 | 38 | loam |
| 1-2C | 35 | 20 | 44 | loam |
| 2-1A | 54 | 14 | 30 | sandy loam |
| 2-1B | 66 | 14 | 18 | sandy loam |
| 2-1C | 54 | 16 | 28 | sandy loam |
| 2-2A | 50 | 16 | 32 | loam |
| 2-2B | 58 | 20 | 21 | sandy clay loam |
| 2-2C | 63 | 10 | 26 | sandy loam |
| 3-1A | 67 | 8.5 | 24 | sandy loam |
| 3-1B | 67 | 10 | 22 | sandy loam |
| 3-1C | 65 | 12 | 22 | sandy loam |

**Table 2.** Annual precipitation (mm) for the three Pinedale, WY area ranches over the course of the experiment, 2012-2015. Precipitation values were derived using interpolated values from the PRISM Climate Group, Oregon State University, http://prism.oregonstate.edu, created 13 Feb 2019.

| Ranch | precipitation (mm) |  |  |  |
| --- | --- | --- | --- | --- |
|  | 2012 | 2013 | 2014 | 2015 |
| 1 | 174 | 257 | 381 | 286 |
| 2 | 209 | 295 | 423 | 354 |
| 3 | 185 | 253 | 378 | 294 |

We used NRCS Soil Survey data to identify five site locations across three privately owned cattle ranches, all within 25 km of the Pinedale Anticline and 10-20 km of each other. Sites were distributed to represent the spatial extent of the proposed mitigation and fertilization treatments (geographic coordinates for specific sites are not provided due to privacy concerns). We obtained permission from three local land-owners to use their privately-owned lands because of constraints associated with imposing manipulations on public areas adjacent to the Pinedale Anticline. We identified site locations with similar soil types, textures, and landscape positions, with all sites positioned on a slight slope. Each site had previously been moderately grazed for over a century; this grazing history is typical of the region, big sagebrush plant communities throughout the western U.S., and - most importantly – consistent with the history of the Pinedale Anticline, for which we are trying to characterize potential effects of N fertilization.

### Experimental design and nitrogen fertilization

To address our hypotheses, we applied three treatments (nitrogen fertilization, nitrogen removal through immobilization, and a no-addition control) in a generalized randomized complete block design with split-plots. Treatments were randomly applied to subplots and replicated in three main plots; this split-plot setup was replicated at five sites (blocks). The hierarchy of locations is site > main-plot > subplot. The five replicate sites were located on three privately-owned ranches. Within each site, we created three plots (10m x 10m). Each main-plot was enclosed by cattle-proof fencing and contained three subplots (2.5m x 8.5m) each randomly assigned one of the above treatments. Subplots were fertilized with N in the form of urea (4.5g N m^−2^) and surface hay (well-aged for at least one year) was applied (referred as ‘nitrogen removal’ here; 250g hay m^−2^) for the above respective treatments in September 2012. The addition of hay to the surface immobilizes soil inorganic N, through incorporation of N into microbial biomass during decomposition of the hay (39–41). All subplots had a 0.56 - 0.625m buffer between subplots and between subplot edge and fence.

### Mineralized Soil Nitrogen

Inorganic nitrogen availability, in the form of nitrate (NO_3_^−^), and ammonium (NH_4_^+^), were estimated *in situ* using Plant Root Simulator (PRS®) probes. The PRS® probes are ion exchange resin membranes held within a rigid plastic frame (Western Ag Innovations, Inc., Saskatoon, Canada) and provide an index of nitrogen availability to plants. PRS probes provide an index of soil N availability that integrates across the time of their deployment (i.e. soil N availability from Oct-May for this study). PRS probes are placed in pairs, one anion probe (collecting NH_4_^+^) and one cation probe (collecting NO_3_^−^). Multiple pairs were placed within a single treatment. All probes were washed with DI water following the manufacturer’s protocol and shipped to Western Ag Innovations, Inc., (Saskatoon, Canada) for analysis.

In order to estimate available soil nitrogen (N) during the first year of the experiment (Sept 2012 – May 2013), we placed PRS® probes within each treatment block at all fifteen plots. We installed probes over winter to catch the spring snowmelt, the largest pulse of nitrogen for the year (42), as plots are inaccessible from winter until soils dry out in the spring.

During the first year they were placed on the edge (i.e., cusp) of the canopy of a big sagebrush individual. During the third year of the study (Oct 2014 – May 2015), probes were placed at two locations to measure soil N available under both the big sagebrush canopy and within the interspace between individual shrubs. For example, if an individual sagebrush had a canopy that extended 20cm from its main stem (i.e., trunk-like base) then the Year 1 probes would be placed at 20cm (i.e., “cusp”), while the Year 3 probes would be placed at 10cm (i.e., “under”) and 30cm (i.e., “interspace” or 10cm outside the canopy) from the stem. Probe placement was stratified by shrub location in an effort to represent a key component of soil N variability in drylands, as N availability is known to be elevated under shrub canopies (43). This was of particular concern as time progressed and the effects of our pulse of N fertilizer on soil N were likely to decrease.

The sets of probes from each sampling campaign (i.e., Year 1 and Year 3) were retrieved from the field and analyzed after their 7-8 month deployment. A limitation of PRS probes is that temperature and soil moisture affect the ability of the anion and cation resin to adsorb NH_4_^+^ and NO_3_^−^. Thus, PRS probes provide an index of N availability that can be compared among treatments within a sampling campaign, but cannot be compared between sampling campaigns. As such, our analyses are limited to comparisons within a campaign (i.e., within Year 1 or within Year 3).

### Sagebrush Growth & Forage Quality

We estimated big sagebrush annual growth by measuring leader growth (i.e., big sagebrush stems) and the ephemeral branch growth (i.e., inflorescences) in 2014. Leaders and ephemeral branches were collected at the end of the growing season in late October 2014 and dried in the laboratory for five days at 55°C and then at 35°C until mass remained constant over a 5-day period. We then weighed and measured the length of each individual leader and branch. The dried leaders were ground in a ball mill and submitted to the Stable Isotope Facility at the University of Wyoming for total carbon and nitrogen analysis (44), as a proxy for forage quality.

Plant Community

We sampled the plant communities at each site over two days in August 2014. We surveyed the plant community using 20 randomly placed Daubenmire quadrats (0.1 m^2^) within each subplot. The canopy cover of each plant species was estimated using a modified Daubenmire (45) method with the following cover categories: <1%, 1-5%, 6-15%, 26-40%, 41- 60%, and >60%. Unknown species were marked as unknown in the field and identified using the Rocky Mountain Herbarium (Univ. of Wyoming). If the unknown species were unable to be identified they were not included in the analyses of the plant community.

### Soil Trace Gas Emission

We sampled soil trace gas emissions twice in July 2013; each was at least 48 hours after any precipitation because high amounts of water vapor can be can confound measurement of CO_2_. All sites were sampled within a single day for each sampling period. We deployed 90 soil gas flux chambers, 2 per subplot across all 15 sites, and arranged them on the cusp of a random interspace and adjacent big sagebrush canopy, as it is well-recognized that a shrub’s location can create important environmental microsites in mountain big sagebrush ecosystems (43).

Sections of polyvinyl chloride (PVC) pipe (10 cm high x 20 cm diameter) were permanently buried in the ground to a depth of 6 ± 2cm, which then acted as bases for the chamber lid. Chamber lids were round, 20cm inside diameter and 8cm deep, made of opaque PVC and vented following Hutchinson & Mosier (46). The chamber base and lid fit snugly together and were sealed using a rubber gasket for the 30-minute sampling period. Once the chamber was sealed, 30 ml samples of air were drawn from the chamber headspace every 15 minutes (t = 0, 15, and 30 mins). Each sample was collected by drawing 60 ml of air from the headspace into a 60 ml syringe (Allison Medical, Inc.), expelling 30ml into the air outside the chamber, and then injecting as much as possible of the remaining 30ml sample into a 12ml flat bottomed, pre-evacuated glass vial (Exetainer, Labco Ltd.) sealed with a rubber septum.

Ambient air samples were collected at each plot following the same procedure above and stored with the sample vials to account for any systemic change during the sample period. Standards of known chemical composition were also carried along on field sampling campaigns and analyzed along with the field samples to account for any systemic change or contamination. Gas samples were analyzed in the laboratory using gas chromatography on a Shimadzu GC-2014 to measure concentrations of CO_2_, CH_4_, and N_2_O.

### Data Analysis

We conducted all data analysis and figure creation using R version 3.5.3 (R core Team, 2018), using the following libraries: knitr, tidyverse, lubridate, plotrix, emmeans, ecodist, labdsv, nlme, Rmisc, vegan, soiltexture, and here.

We used a linear mixed effects model (*lme* function, ‘nlme’ package v 3.1-139 (47)) to examine the fixed effects of treatment with the nested, random effects of site and plot (i.e., y ~ treatment + 1 | site / plot). Treatment effects were first evaluated using a one-way analysis of variance (ANOVA), followed by an examination of contrasts to determine the effectiveness of individual N and hay treatments. Finally, we conducted a post-hoc evaluation of the 95% confidence intervals for the response to each treatment using estimated marginal means (*emmeans* function, ‘emmeans’ package v1.3.4 (48)).

The mixed effects model was used to evaluate the effects of our N and hay treatments on inorganic soil N availability (total inorganic N, NO_3_^−^, NH_4_^+^), sagebrush growth (leader length, inflorescence length, leader dry mass, and leader dry density), and forage quality (percent C, percent N, C:N ratio, leader N mass, leader C mass).

Soil trace gas fluxes (CO_2_, CH_4_, N_2_O) were calculated for each treatment and plot using the rate of change for each of the gases analyzed over the 30-minute period, calculated as the slope values of a linear regression. These fluxes were then similarly analyzed using the mixed effects model previously described.

We examined the effects of N and hay treatments on the plant community by evaluating differences in total plant cover and species richness using the same mixed effects model and associated processes. To explore the effects of treatment applications on plant community composition, we used a permutational multivariate analysis of variance (PERMANOVA). Total cover and abundance by species was assessed by summing the cover class midpoint for each species by site to obtain the total percent cover for each species and then standardized these values to relative abundance (*decostand* function, ‘Vegan’ package v2.5-4 for R (49)). We then used a PERMANOVA to evaluate differences in community composition between treatments using a Bray-Curtis dissimilarity index and 9999 permutations (*adonis* function, ‘Vegan’ package v2.5-4 for R).

We also explored the effects of total soil N on leader length through calculating a one-way analysis of covariance (ANCOVA) with site and plot as nested, random factors. This is similar to the original model described above except treatment is removed, as it is confounded with soil N availability. We then obtained fitted values at the population and hierarchical level (i.e., individual sites).

## RESULTS

### Total Inorganic N in Soil

The addition of N significantly increased the availability of total inorganic N in soils (p<0.001; Fig 1) during the first year of the experiment, while hay addition significantly reduced the availability of N (p=0.035; Fig 1). No differences were found for availability of NH_4_^+^ between treatments, however NO_3_^−^ was significantly more available in the N fertilized treatments (p<0.001) than either the control or N removal (p=0.035). By the end of second year of the experiment, no significant differences remained between treatments for total inorganic N, NO_3_^−^, or NH_4_^+^. During year one, NO_3_^−^ represented 99.0% ± 0.4 of the total available inorganic N across all treatments/locations, while in year two it was 92.0% ±1.7.

**Fig 1.**
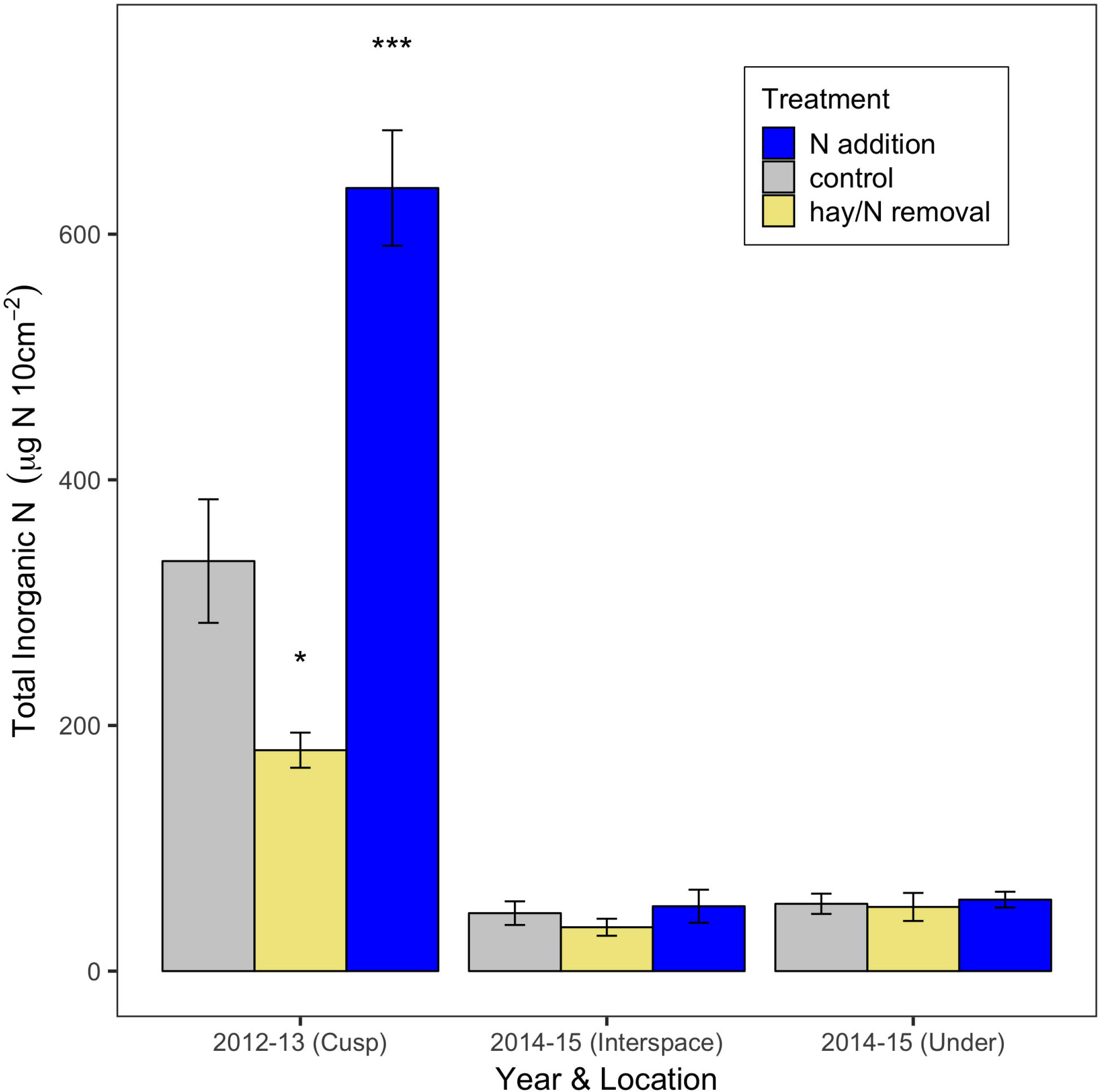
Mean total soil inorganic N availability within a big sagebrush plant community in Pinedale, WY. N fertilization occurred in 2012 with soil available inorganic N measured via PRS probes in fall/winter/spring of 2012-2013 and 2014-2015 (n=5 sites). The 2013 data were collected from soil on the *cusp* of the canopy and interspace, while 2015 data were collected separately from within the *interspace* and *under the canopy*. Significant differences from the control, within each year and location, are indicated by asterisk based on α = 0.05 (* indicates p<0.05, *** indicates p<0.001). Whiskers indicate the standard error based on among-site variation.

### Leader Growth & Nitrogen Content

The addition of N and the application of hay had no treatment effect on the growth of big sagebrush leaders; they were unchanged in length, mass, and density. While the treatment term was significant in predicting leader length (p=0.006), neither the nitrogen additions (p=0.103) nor hay treatment (p=0.079) were significantly different than the control when examining contrasts. Ephemeral branches (i.e., inflorescences) were unchanged by the addition of N fertilizer, though the addition of hay reduced their mean length by approximately 8% or 1.1cm (p=0.029; Fig 2). Leader C:N ratio and N content (i.e., forage quality) were also unchanged by treatment application.

**Fig 2.**
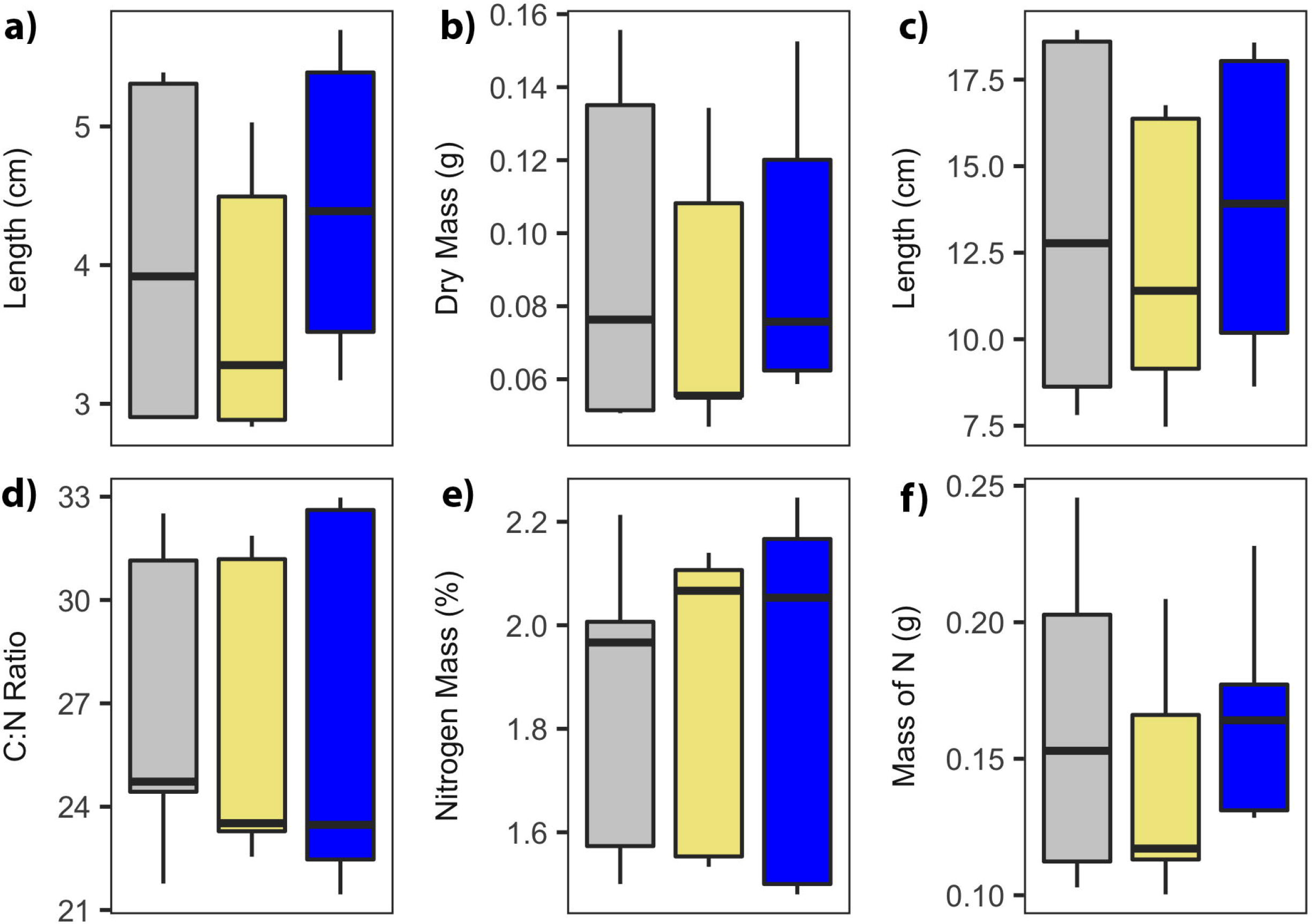
Growth of sagebrush leaders was unaffected by treatment application, while inflorescences decreased in length with hay addition. Treatment is indicated by color (gray=control, khaki=hay addition, blue=nitrogen addition). a) Leader length, b) leader dry mass, c) inflorescence length, d) leader C:N ratio, e) leader N content in percent, and f) leader N content in grams.

Exploring the relationship between soil total N availability (Fig 3), we found that the coefficient for soil N was significantly different from zero (slope=0.00122 ± 0.0003259 SE, t=3.74365, p=0.0008).

**Fig 3.**
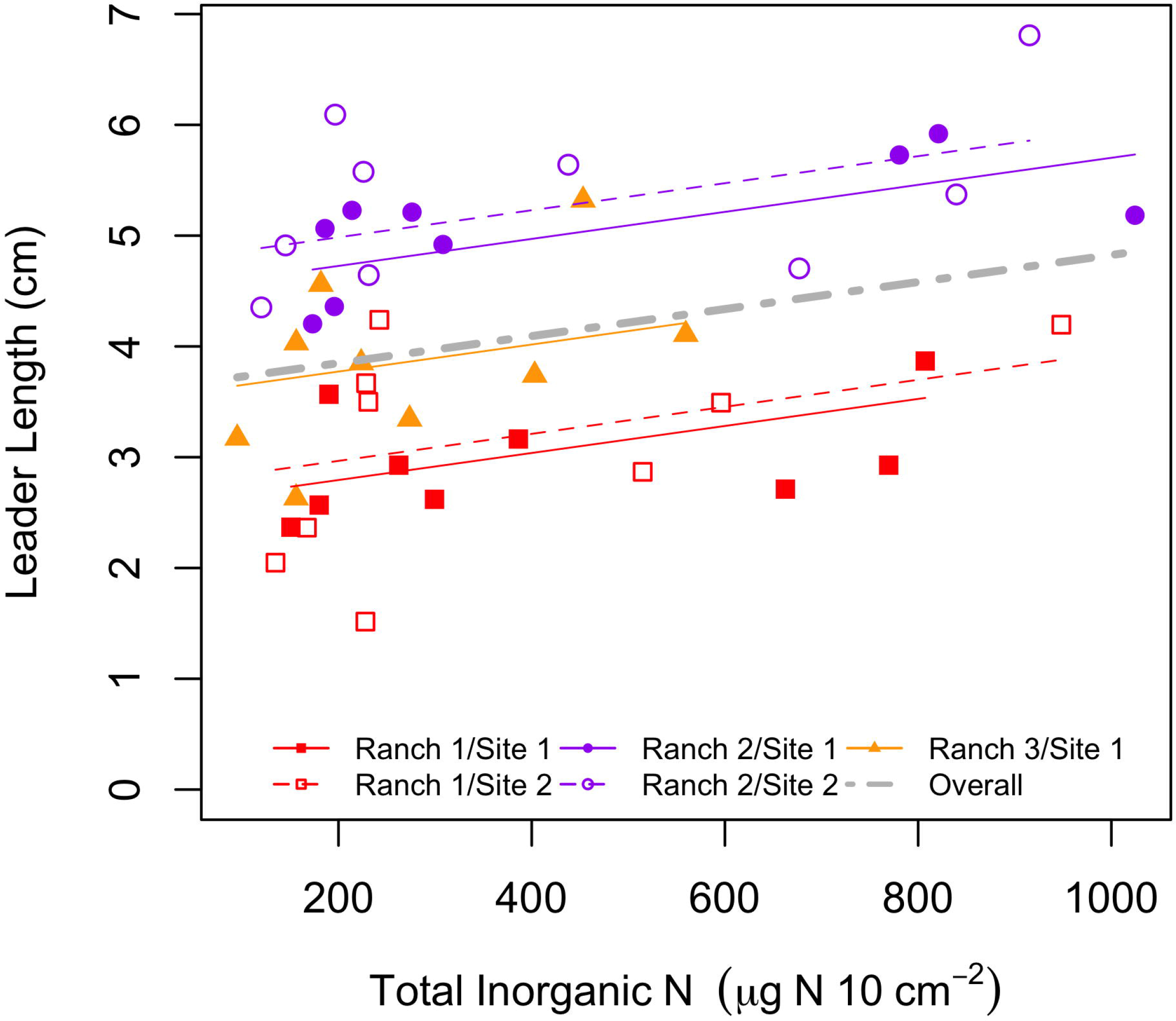
Relationship between sagebrush leader growth and total inorganic N for each of five sites in Pinedale, WY. Ranch location is indicated by color, while sites within a ranch differ in shape (note: Ranch 3 has only a single site). Solid lines correspond to the site with the solid color shapes and dashed lines correspond to the hollow/un-filled shapes. Data presented is across all treatments.

### Plant Community

The three ranches each had similar big sagebrush density, typically ranging from 20-30% of cover. The most common understory species, by cover, across the controls plots within the three ranches included *Chrysothamnus viscidiflorus*, *Pascopyrum smithii*, *Poa secunda*, *Achnatherum pinetorum*, *Hesperostipa comata*, and *Phlox hoodii*. All common understory species were found at each site, though their prevalence did vary slightly between ranches.

We found no treatment effect on total plant cover or species richness (Fig 4). The plant community also remained unchanged, regardless of the treatment application.

**Fig 4.**
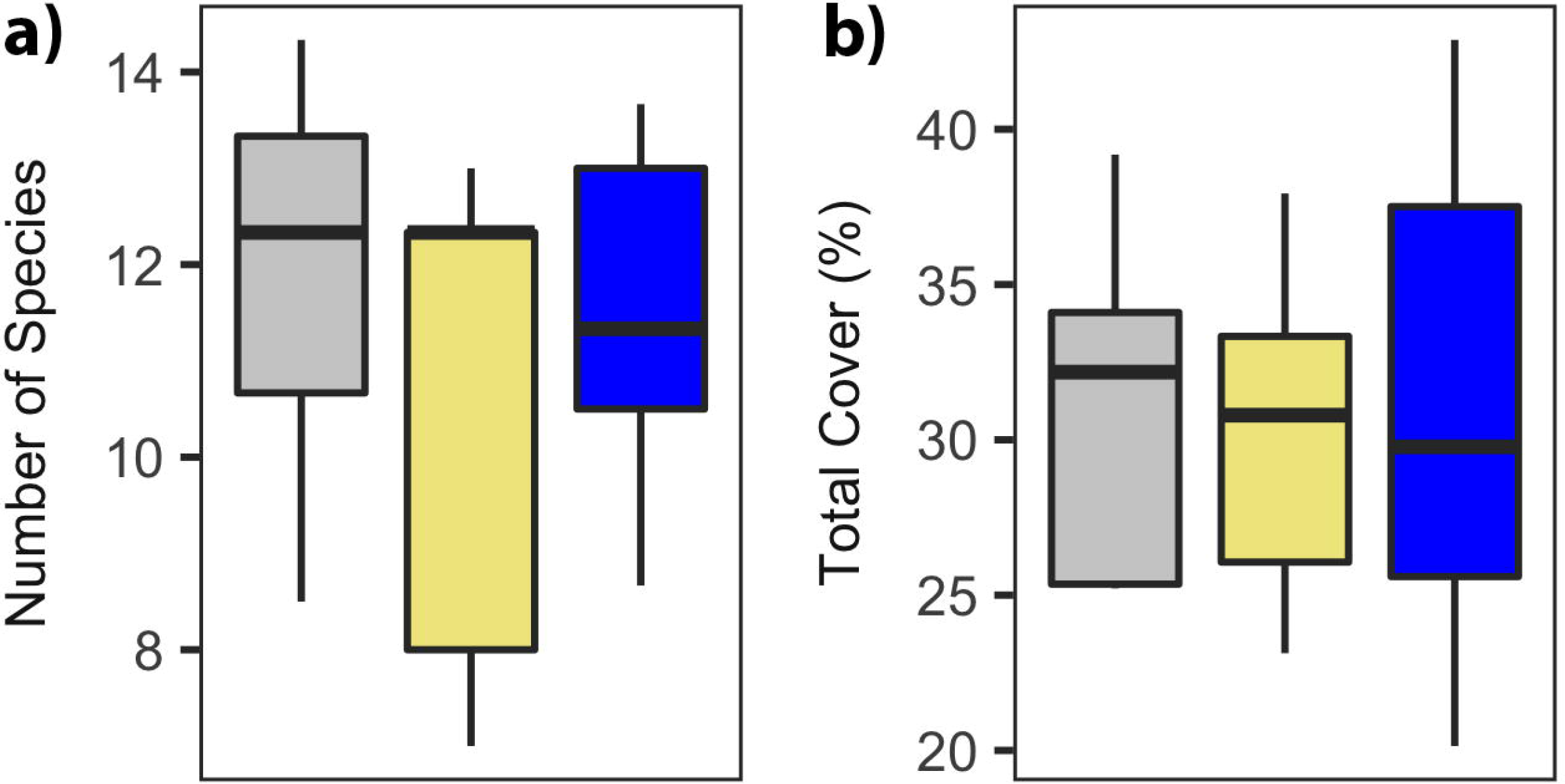
Total plant cover and species richness were unaffected by treatment application. Treatment is indicated by color (gray=control, khaki=hay addition, blue=nitrogen addition). a) species richness, and b) plant abundance

### Trace Gas Emission from Soil

Soil trace gas emission displayed no effect from treatments for any gas species (Fig 5). Neither the addition of hay or N altered the efflux of CO_2_, CH_4_, or N_2_O from the soil surface.

**Fig 5.**
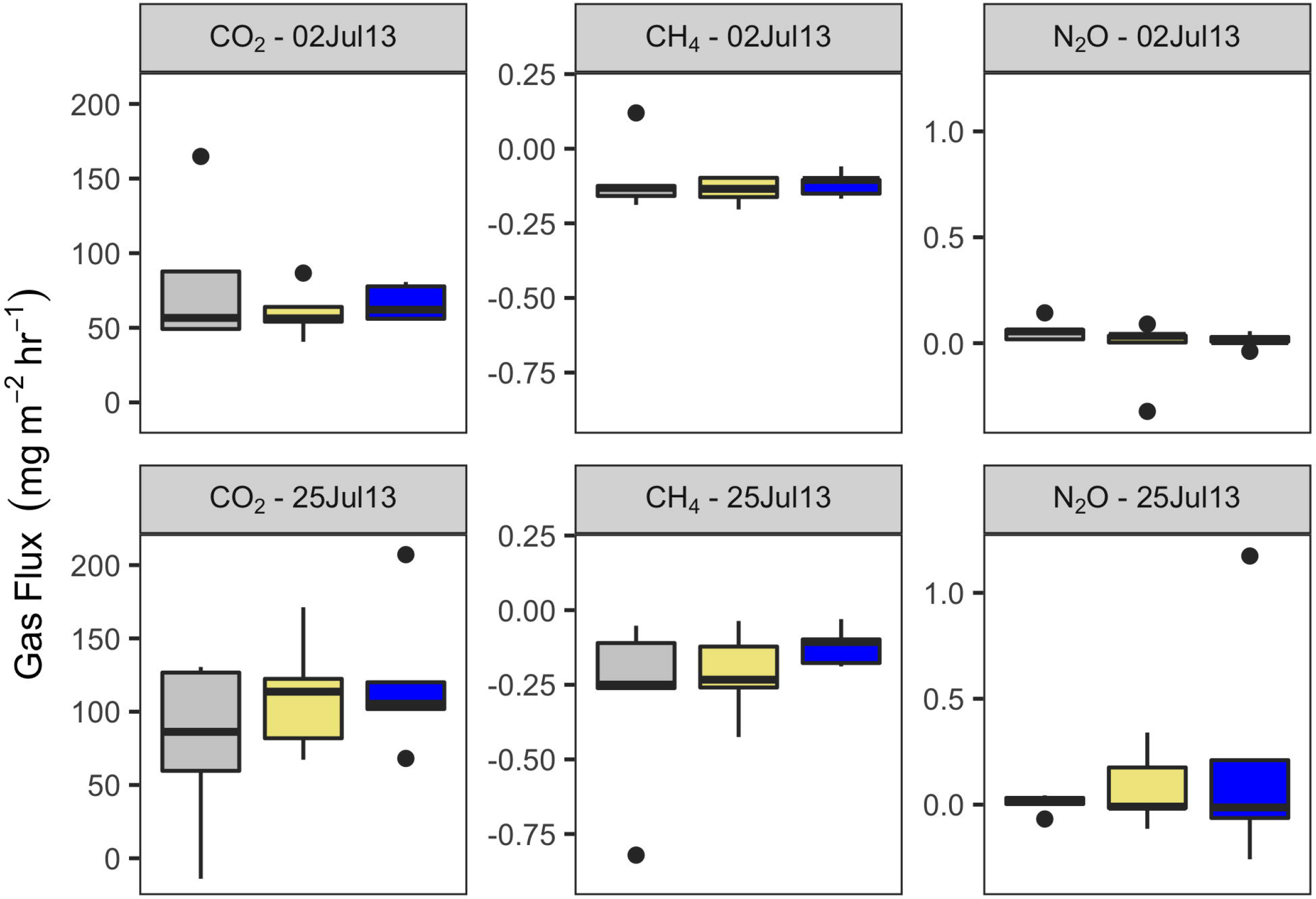
Trace gas emissions (CO_2_, CH_4_, & N_2_O) were unaffected by treatment application. Treatment is indicated by color (gray=control, khaki=hay addition, blue=nitrogen addition).

## DISCUSSION

N fertilization significantly increased the availability of N in soil for the first-year post-treatment, however there was no significant difference in N availability by the second year. Differences in control values between year one and two are not surprising given the different locations; length of deployment, soil temperature, and soil moisture can all affect the adsorption of anion and cations to their counter-ion resins, making comparisons between years (and conditions) inappropriate to a certain extent. Precipitation between the two years, 2012 and 2014, varied by approximately 200mm depending on the ranch examined and could account for a portion of the variability seen in the control plots between years. Due to differing conditions between years, we have focused on comparing PRS probe measurements of total inorganic N within single years.

The lack of significant increase in soil N availability in the second year in fertilized plots is likely due to ‘normal’ N losses through N mineralization, gaseous loses (volatilization and microbially-mediated), and leaching. Additions of N were conducted at 4.5 g N m^−2^ (or 45 kg N ha^−1^), which is 150-900% of annual N mineralization (43). Trace gas emissions of N-containing gases (N_2_O and NO_x_) accounted for an additional 10-20 kg N ha^−1^ yr^−1^ loss from the soil, which could contribute to the lack of increase observed in inorganic soil N among treatments in year two (5,26). While we found lower fluxes of N_2_O compared to these studies one year after fertilization, it may be that fluxes were greater immediately after fertilization or during the spring when water was more available, therefore not captured in our sampling period.

Increased N availability had minimal effect on the length of big sagebrush leaders over the two years of the study. Additional N did not result in an increase in forage quality; C:N ratio and mass of N were the same between the control and N fertilized plots. While the addition of hay minimally decreased the length of ephemeral branches, the addition of N had no effect. It is possible that the decrease in soil N caused by the hay treatments decreased growth by increasing N limitations. These limitations on the growth of inflorescences by N may have been more apparent during 2014, when water was more available.

Our prediction that there would be no differences in plant growth between treatments was generally supported. Conversely, we also found that leader length was weakly positively correlated with inorganic soil N availability at each of our sites. On the face of it, these two statements are contradictory. While we have shown that our treatments significantly affected the availability of total N in soil across all sites, there was no associated treatment effect on plant growth. Leader growth correlated with N availability at the site level. It is possible that other factors may drive or mediate plant growth at these sites. Site-specific factors such as soil texture (i.e., clay), precipitation, and leaching can affect the conversion of urea fertilizer to N, the retention of that in soil, and/or to the uptake and allocation of that N by plants to increased leader growth.

Interestingly, differences in leader length between ranches was strong. While this could be due to many factors, the ranch with highest precipitation in each year from 2012-2015 also had the longest sagebrush leaders. However, we did not measure precipitation at any of our sites or ranches, making it impossible to assert this with any confidence.

Contrary to our predictions, the addition of a single pulse of N did not alter plant community composition. *P. smithii* (western wheatgrass) is known to be nitrophilic and respond to additions of N (50,51), however no significant change was observed. In addition, treatments did not affect either total plant cover or species richness. This is not surprising as soil N availability was only increased for the first year after treatment. In addition, drylands – and big sagebrush ecosystems – are primarily water limited, making short-lived increases of N unlikely to affect cover or richness.

There was no effect of N additions on GHG efflux (i.e., CO_2_, CH_4_, and N_2_O) from soil. Fluxes of CO_2_ and N_2_O fell into similar ranges for dry, unwetted sagebrush reported by Norton et al. (52). The lack of significant treatment effect could be due to a number of reasons: (1) trace gas measurements are highly variable and sensitive to a number of different environmental factors; (2) increased GHG fluxes occurred during a time that was not measured, potentially closer to fertilization or during the spring when water is more available; or (3) the ability of the microbial community to produce CO_2_, CH_4_, and N_2_O may be limited primarily by water availability, not N availability.

Our findings are not consistent with other studies, which have generally found a positive response of plant productivity to N fertilization (1,8) indicating some N limitation. However given that the mean annual precipitation of the Pinedale, WY area is 290mm, it is not surprising that we found no response of leader growth to N additions; Hooper and Johnson (1) found that the response to N was greater for areas with 450-900 mm of precipitation than for 300 mm. It is important to note that while most studies find a primary water limitation in drylands (53), the response of dryland ecosystems to N can be expected to increase with increased precipitation (54).

Our results indicate that a one-time application or single pulse of N does not affect soil N availability in the long term within the sagebrush communities of the Pinedale Anticline. We suggest that annual fertilization would be necessary to maintain high levels of soil N availability within big sagebrush systems, as the levels of soil N were the same between the fertilized and control plots by the second year. Yet increases in N availability over the long term may have negative consequences, including higher volatile N losses, and increases in invasive plants (26). It should be noted that the chronic additions of N fertilizer would likely have different effects than that of the single pulse of N applied during this study. This study has addressed the effects of a single pulse of N, such as the PAPO pilot study on the Pinedale Anticline. The increased levels of N found in the first year after fertilization only minimally increased the length of big sagebrush leaders, and continued application at that level would be unlikely to increase forage quality. Ecosystem processes in drylands are primarily water limited and do not change with short-term N additions, at least over the two-year duration of this study.

## CONCLUSIONS

This study was conducted to investigate the role of N limitation in dryland sagebrush ecosystems and to consider the effect of a pulse of N addition to increase forage and forage quality for mule deer. We evaluated responses of soil N availability, annual growth rate of big sagebrush, and forage N concentration in response to N additions. In addition, we investigated the response of trace gas production and plant community structure to increased N. This study demonstrates that a single addition of N, in the form of urea fertilizer, does increase N availability in soil for one year after fertilization. However, this increased soil N availability is only loosely associated with increased growth of big sagebrush and not likely to impact forage availability and quality for mule deer. The results of this study suggest that the addition of N in a single pulse is likely a poor choice for mitigation of mule deer population declines on the Pinedale Anticline due to the minimal gains in forage, if any, and associated high cost.

## ACKNOWLEDGEMENTS

Thank you to B. Amgalan, O. Avirmed, D. Bell, M. Cleary, T. Martyn, K. Palmquist, V. Pennington, B. Peterson, and C. Rottler for assistance with field work, data collection, lab work, and manuscript review. Additional thanks to D. Schlaepfer and T. Gregoire for help with data analysis. Thank you to our three private landowners for making this research possible, including Freddie Botur and the Cottonwood Ranch in Big Piney, as well as the Sommers Ranch in Pinedale. An earlier poster version of this study was presented at the Ecological Society of America conference in 2015. Many thanks to those attendees that provided feedback and encouragement. Financial support for this research came from the University of Wyoming and the Wyoming Excellence Fund.

